# Mosquito Tissue Ultrastructure-Expansion Microscopy (MoTissU-ExM) enables ultrastructural and anatomical analysis of malaria parasites and their mosquito

**DOI:** 10.1101/2024.04.17.589980

**Authors:** Benjamin Liffner, Thiago Luiz Alves e Silva, Joel Vega-Rodriguez, Sabrina Absalon

## Abstract

Study of malaria parasite cell biology is challenged by their small size, which can make visualisation of individual organelles difficult or impossible using conventional light microscopy. In recent years, the field has attempted to overcome this challenge through the application of ultrastructure expansion microscopy (U-ExM), which physically expands a biological sample approximately 4.5-fold. To date, U-ExM has mostly been used to visualise blood-stage parasites and used exclusively on parasites *in vitro*. Here we develop <u>Mo</u>squito <u>Tiss</u>ue <u>U-ExM</u> (MoTissU-ExM), a method for preparing dissected mosquito salivary glands and midguts by U-ExM. MoTissU-ExM preserves both host and parasite ultrastructure, enabling visualisation of oocysts and sporozoites *in situ*. We validate that MoTissU-ExM samples expand as expected, provide a direct comparison of the same dissected tissues before and after MoTissU-ExM, and highlight some of the key host and parasite structures that can be visualised following MoTissU-ExM. Finally, we provide a point-by-point protocol for how to perform MoTissU-ExM, along with details on how best to image the expanded tissues, and how to troubleshoot common issues.

## Introduction

Cell biology is fundamentally underpinned by our ability to visualise cells and use this information to infer biological structure and function. Light microscopy, the visualisation of cells based on the light they emit or absorb, is one of the most widely used tools in cell biology. Historically, the ability of light microscopy to resolve biological structures had been bounded by diffraction limit of light^1^. Over the last few decades, however, many methods that overcome this limit, collectively called “super-resolution” microscopy, have been developed. Typically, super-resolution microscopy techniques involve the use of some form of specialised instrumentation or analysis to improve resolution and therefore the ability to visualise biological structures^2^. In 2015, however, a conceptually different method called expansion microscopy (ExM) was developed^3^. Rather than use specialised instrumentation, ExM is a sample preparation method that physically expands a biological sample; resulting in a dramatic increase in the ability to resolve biological structures without the need for specialised equipment.

Since the initial development of ExM, many adaptations to and variations of ExM have been published. Notably, ultrastructure expansion microscopy (U-ExM)^4^ has been widely applied to the study of single-celled organisms^5-12^. U-ExM involves the tethering of a biological sample to a hydrogel using formaldehyde and acrylamide, before denaturation, expansion and staining. Importantly for its use in single-celled organisms, U-ExM results in near-native preservation of cellular ultrastructure and 4 to 4.5-fold isotropic expansion^4^.

U-ExM has been widely adopted in the study of the cell biology of apicomplexan parasites^13^. Apicomplexa are a phylum of largely parasitic single-celled organisms that include significant causes ofdisease in humans and livestock, such as malaria, toxoplasmosis and cryptosporidiosis. To date, U-ExM has been used to study the cell biology of *Plasmodium*^6,14,15^, *Toxoplasma*^16,17^, *Cryptosporidium*^7,18^, and *Neospora*^19^. To date, U-ExM has only been applied to parasites cultured *in vitro* and prepared as either isolated parasites or infected-cell monolayers. Being able to observe the anatomical context these parasites exist in, however, is key to understanding both parasite biology and host-parasite interactions. Here, we develop a method we call <u>Mo</u>squito <u>Tis</u>sue <u>U-ExM</u> (MoTissU-ExM), a protocol tailored to visualise the ultrastructure of malaria parasites (*Plasmodium*) and their mosquito hosts *in situ*. We validate that this method fully expands the mosquito tissues and we test a range of commonly used fluorescent dyes and stains. Finally, we provide a point-by-point protocol for dissecting, preparing, and imaging expanded mosquito tissues. While developed for malaria parasites, the application of MoTissU-ExM could extend across various fields of entomological and parasitological research.

## Results

U-ExM has been applied to different stages of the malaria parasite lifecycle but not to either mosquito midgut oocysts or salivary gland sporozoites. Whole tissues have previously been visualised using U-ExM^20^, including zebrafish and mouse embryos, and *Drosophila* wings, but this involved modifications to the U-ExM protocol that more than doubled sample processing time. We reasoned that dissected mosquito midguts and salivary glands are enclosed epithelial monolayers and, therefore, would not require the extensive processing required for thicker tissues.

### Development of Mosquito Tissue U-ExM (MoTissU-ExM)

We then developed a pipeline for the dissection, fixation, gelation, expansion, and imaging of both infected mosquito midguts and salivary glands (Figure 1) (see point-by-point protocol). Briefly, dissected mosquito tissues were fixed in 4% v/v paraformaldehyde before in-solution anchoring in formaldehyde/acrylamide. Anchored tissues were concentrated and transferred to 12 mm Ø poly-D-lysine-coated coverslips (approx. 5 tissues per coverslip). Excess anchoring solution was removed, tissues were evenly spaced apart, and underwent gelation. Following gelation, gels were placed in denaturation buffer to separate the gel and coverslip. Once gels had separated from the coverslip, each infected tissue (still visible in the gel) was cut out of the gel for individual denaturation, expansion, staining, and imaging.

**Figure 1:**
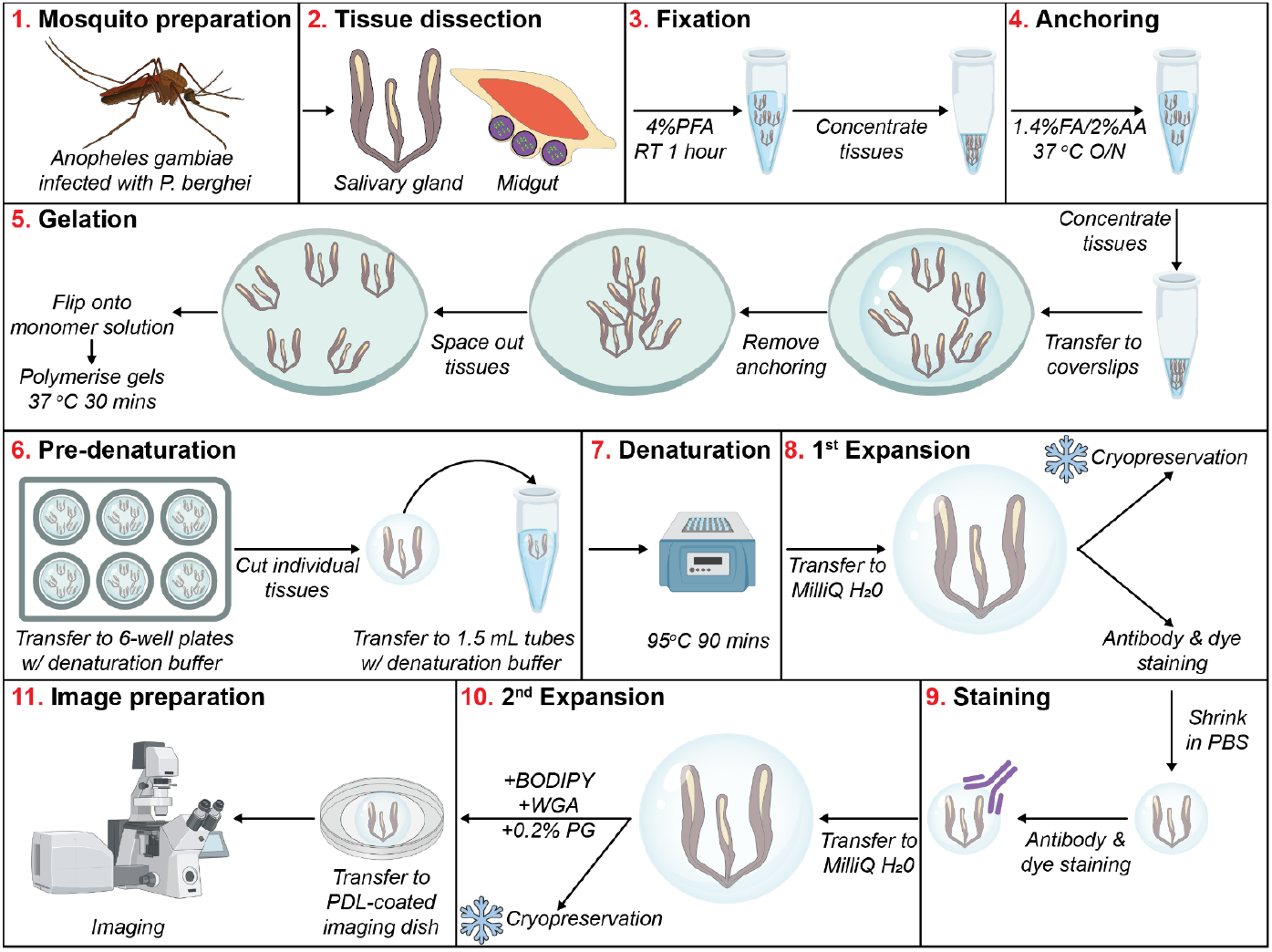
Workflow for preparing <u>Mo</u>squito <u>Tiss</u>ues using <u>U-ExM</u> (MoTissU-ExM). Workflow diagram for the dissection, preparation, and expansion of mosquito tissues. Note that the workflow alternates between being read left to right (steps **1**. – **4**. and **6**. – **8**.) and right to left (steps **5**. And **9**. to **11**.) between rows. A detailed step-by-step text version of this process can be found in the methods section. Steps **3**. – **11**. depict salivary glands, but the process is the same for midguts. Longitudinal opening of midguts is not depicted in this workflow. PFA: paraformaldehyde, RT: room temperature: FA: formaldehyde, AA: acrylamide, O/N: overnight, DI H2O: deionised water, PBS: phosphate buffered saline, WGA: wheat germ agglutinin, PG: propyl gallate. Snowflakes indicate stages where gels can be cryopreserved at -20 °C.

### Dissected mosquito tissues expand fully

When a sample is appropriately denatured in the U-ExM protocol, the diameter of the expanded gel divided by the diameter of the unexpanded gel can be used as a guide for the expansion factor of the biological sample^6^. For this method, mosquito tissues were cut individually from the gel, hence the expansion factor could not be estimated in this way. To determine the expansion factor of mosquito tissues prepared by U-ExM, we measured the average diameter of nuclei in both unexpanded and U-ExM midguts (Figure 2) and salivary glands (Figure 3). Additionally, we measured the same nucleus in a salivary gland (Figure 2a) or a midgut (Figure 3a), both before and after expansion. In unexpanded midguts (Figure 2b), the average midgut epithelial cell nucleus diameter was 7.252 µm (± SD 1.57 µm, 200 nuclei, 4 midguts), while the average nucleus diameter in U-ExM midguts was 31.62 µm (± SD 5.79 µm, 200 nuclei, 4 midguts). In unexpanded salivary glands (Figure 3b), the average salivary gland epithelial cell nucleus diameter was 10.17 µm (± SD 1.66 µm, 179 nuclei, 4 salivary glands), while the average nucleus diameter in U-ExM salivary glands was 42.63 µm (± SD 8.35 µm, 176 nuclei, 5 salivary glands). This estimates an average linear expansion factor of 4.19-fold for salivary glands and 4.36-fold for midguts across multiple experiments. These expansion factors are consistent with those previously published using U-ExM^4,6,14,21^ and suggest that mosquito tissues prepared using U-ExM are fully expanded.

**Figure 2:**
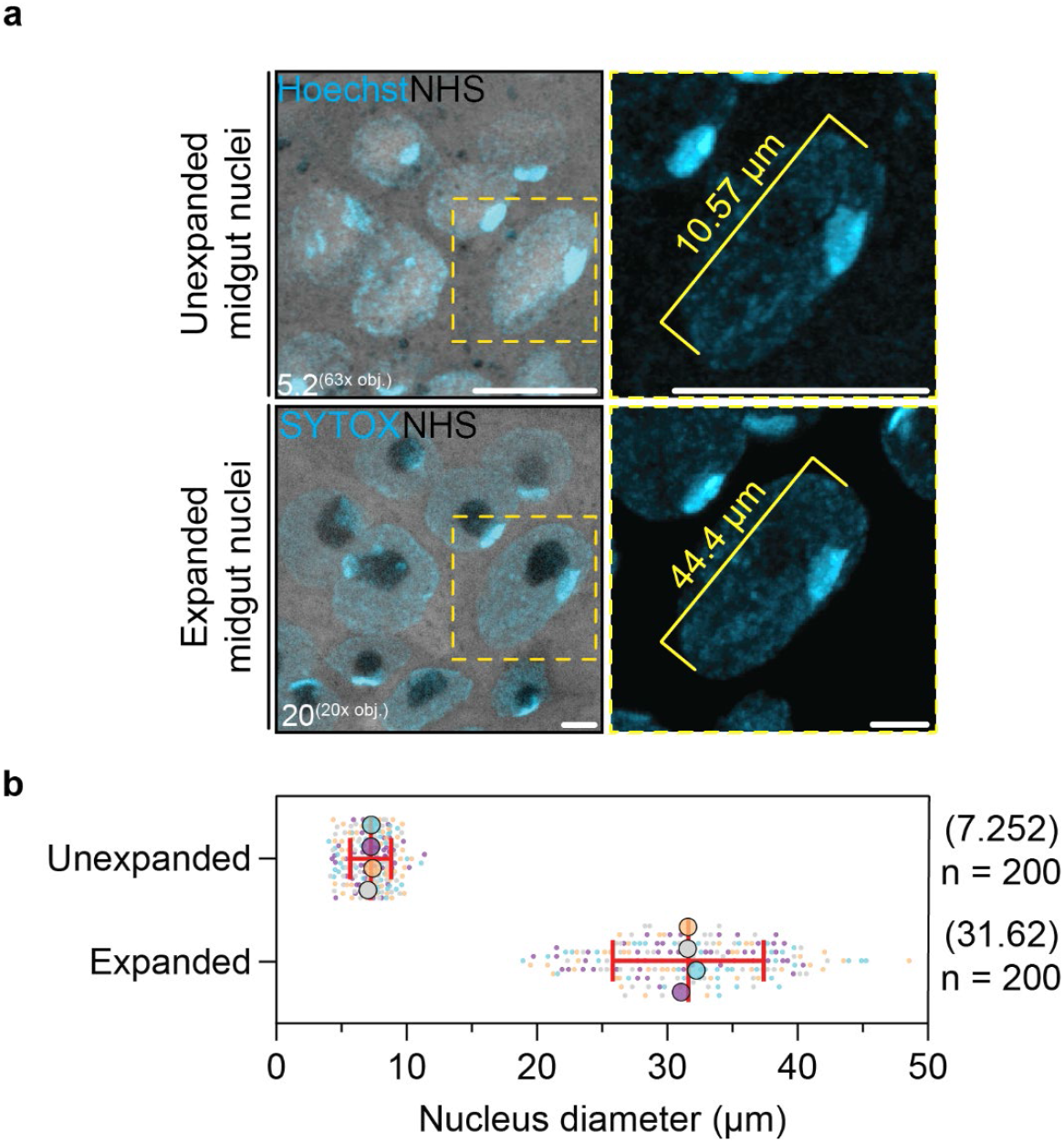
Measuring expansion factor of MoTissU-ExM midguts. **(a)** To determine the expansion factor of midguts prepared by U-ExM, unexpanded midguts were stained with Hoechst (cyan, DNA) and NHS ester Alexa Fluor 405 (greyscale, protein density), and imaged by Airyscan 2 microscopy. U-ExM midguts were stained with SYTOX (cyan, DNA) and NHS ester. The same nucleus, before and after U-ExM is depicted. Number in the bottom corner of each image indicates z-projection depth in µm. obj.= objective lens (see methods section for objective lens details). Scale bar = 10 µm. **(b)** Maximum midgut nucleus diameter was measured in both unexpanded and U-ExM midguts, giving an average estimated expansion factor of 4.36-fold. Approximately 50 nuclei from 4 unexpanded and U-ExM midguts were measured. Small datapoints represent individual nuclei measurements, while large datapoints represent midgut means. Error bars = SD.

**Figure 3:**
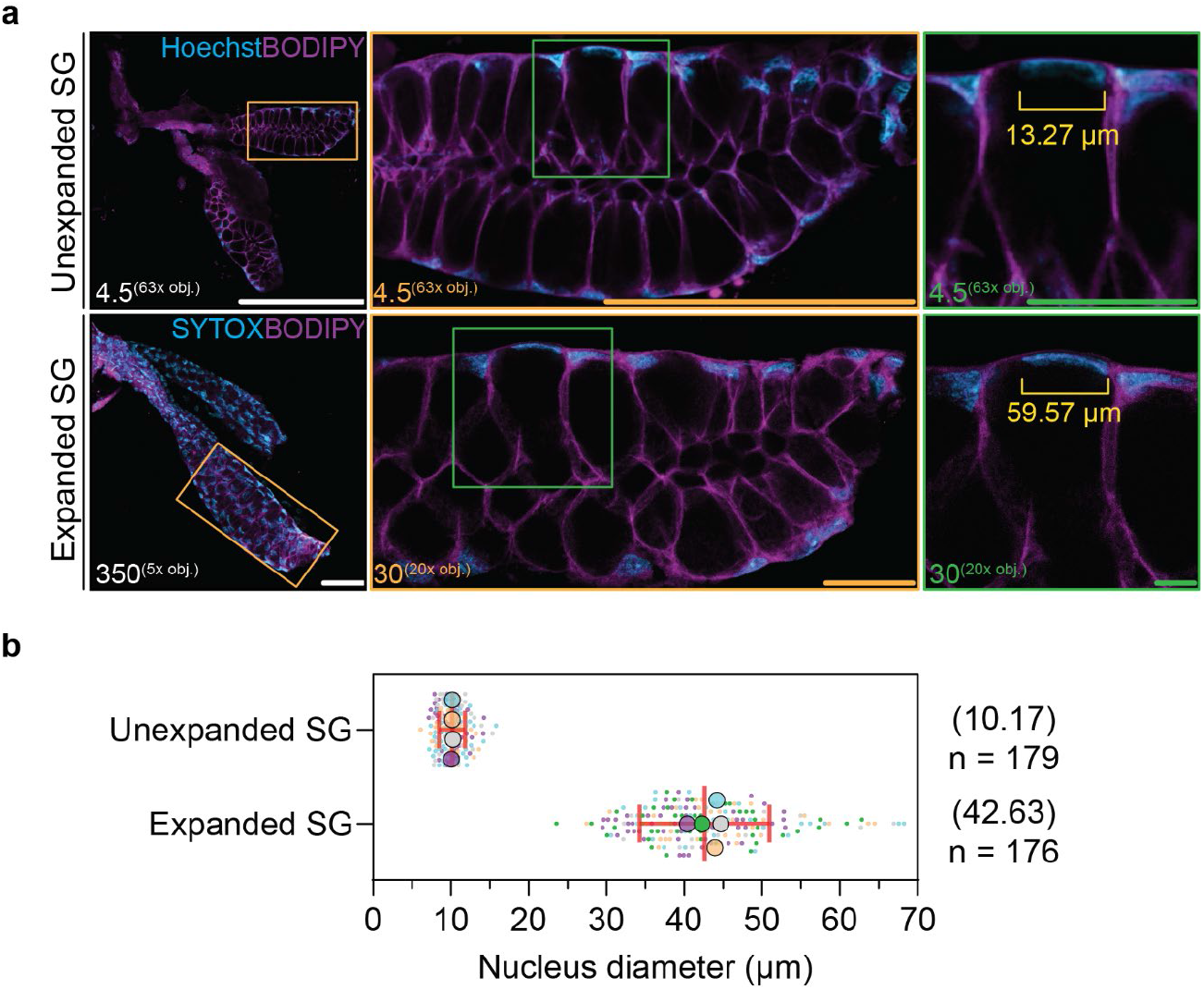
Measuring expansion factor of MoTissU-ExM salivary glands. **(a)** To determine the expansion factor of salivary glands prepared by U-ExM, unexpanded midguts were stained with Hoechst (cyan, DNA) and BODIPY-FL-Ceramide (magenta, lipids), and imaged by Airyscan microscopy. U-ExM salivary glands were stained with SYTOX (cyan, DNA) and BODIPY. The same salivary gland, before and after U-ExM, is depicted (note that salivary gland orientation changed during U-ExM preparation) with a zoom of the same lateral lobe (orange) and same nucleus (green). Number in the bottom corner of each image indicates z-projection depth in µm. obj.= objective lens. Scale bars: = 200 µm, orange = 50 µm, green = 20 µm. **(b)** Maximum salivary gland nucleus diameter was measured in both unexpanded and U-ExM salivary glands, giving an average estimated expansion factor of 4.19-fold. 179 nuclei from 4 unexpanded salivary glands, and 176 nuclei from 5 U-ExM salivary glands were measured. Small datapoints represent individual nuclei measurements, while large datapoints represent salivary gland means. Error bars = SD.

### U-ExM of mosquito tissues preserves both parasite and host ultrastructure

Highly chitinous tissues from arthropods can be resistant to isotropic expansion^22^, introducing sample distortions. Given that nuclei expanded as expected, it was unlikely that chitin was limiting expansion in mosquito salivary glands or midguts, but we wanted to confirm that both the host tissue and the parasites were being preserved at both the anatomical and ultrastructural levels.

In U-ExM midguts, both large anatomical features like muscle fibres and fine anatomical features like microvilli were preserved (Figure 4) in a manner similar to previous electron microscopy studies^23,24^. Additionally, *P. berghei* oocysts were well preserved with the oocyst capsule along with developing sporozoites and their DNA (Figure 4). For a more detailed comparison, we imaged the same midgut, same oocyst, and same forming sporozoites both before (Figure 5a) and after (Figure 5b) expansion. The entire midgut, along with the relative position of all host cells and oocysts within that midgut appeared highly preserved with no significant aberrations observed at either the anatomical or ultrastructural levels.

**Figure 4:**
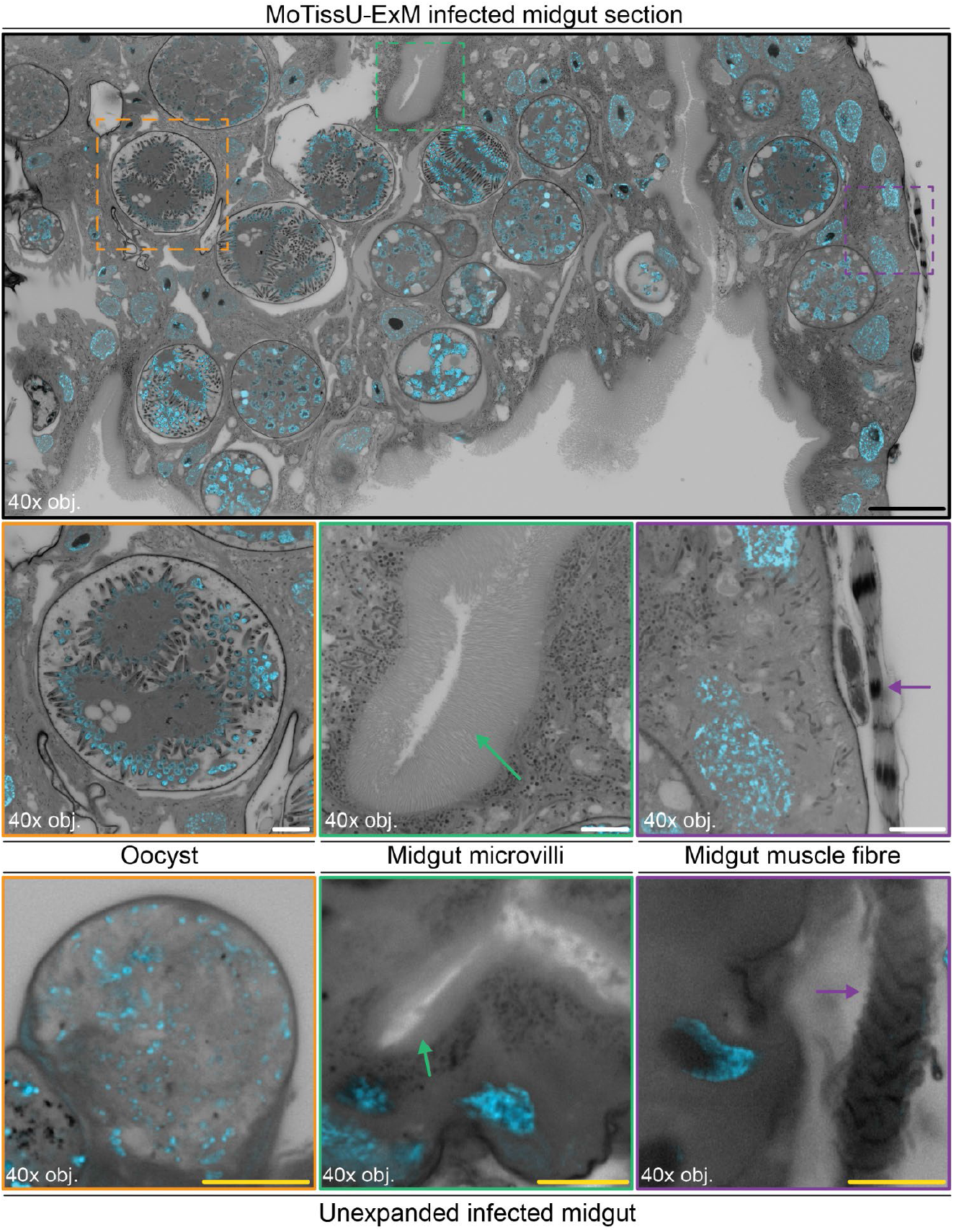
Mosquito midgut U-ExM preserves both parasite and host ultrastructure. *P. berghei* infected mosquito midguts were prepared by U-ExM, or left unexpanded, stained with SYTOX (cyan, DNA) and NHS ester Alexa Fluor 405 (greyscale, protein density), and imaged by Airyscan Microscopy. From a larger section of the mosquito midgut (black), ultrastructural conservation of both parasite (oocyst, orange) and host (microvilli, green arrows & muscle fibres, magenta arrows) can be observed. Note that images of the unexpanded midgut are not the same midgut prepared by U-ExM. No significant gross, or ultrastructural abnormalities were observed in either the mosquito tissue or parasite following U-ExM. obj.= objective lens. Black scale bar = 100 µm, White scale bar = 20 µm, yellow scale bar = 10 µm. All images are a single z-slice.

**Figure 5:**
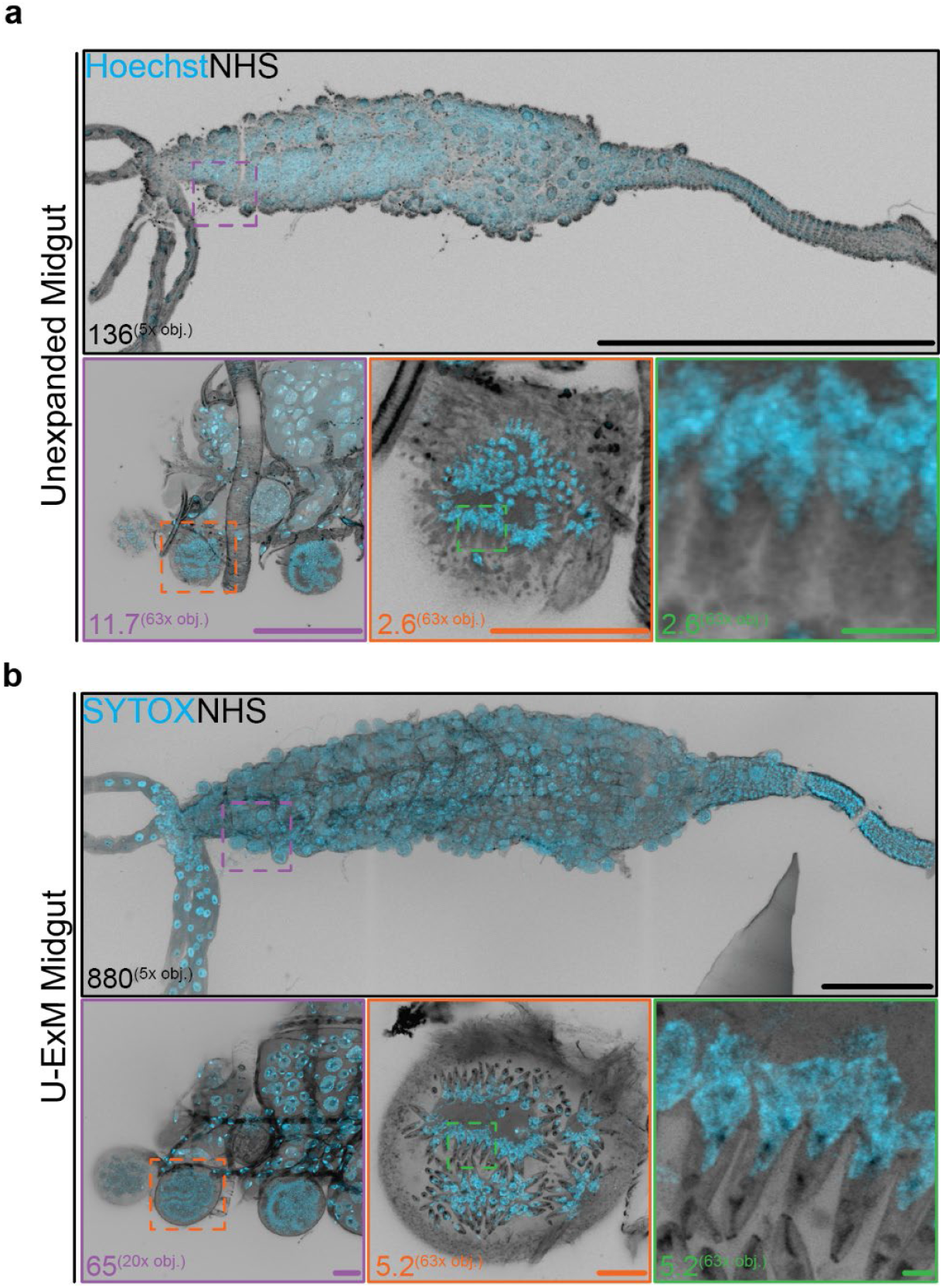
Imaging of an infected midgut before and after MoTissU-ExM. **(a)** A mosquito midgut infected with *P. berghei* was fixed, stained with Hoechst (cyan, DNA) and NHS ester Alexa Fluor 405 (greyscale, protein density), and imaged by Airyscan microscopy. The entire midgut (top row), a section of that midgut (magenta), an individual oocyst (orange), and forming sporozoites in that oocyst (green) are all depicted. **(b)** After imaging, the same midgut, section, and oocyst were prepared by U-ExM, stained with SYTOX (cyan, DNA), re-stained with NHS Ester, and re-imaged. Number in the bottom corner of each image indicates z-projection depth in µm. obj.= objective lens. Scale bars: white black = 1000 µm, magenta = 50 µm, orange = 25 µm, green = 5 µm.

Similarly to midguts, both salivary gland morphology and ultrastructure were preserved following U-ExM. In expanded salivary glands, lateral and medial lobes were easily distinguished from each other, with a clear distinction between the distal and proximal regions of those lobes (Figure 6). Further, throughout the whole salivary gland, the secretory cavity could be easily distinguished from the cytoplasm of the epithelial cells just by using stains for protein density, wheat germ agglutinin (WGA), or lipids (BODIPY) (Figure 6). For sporozoites in expanded salivary glands, their relative position (epithelial cell cytoplasm or secretory cavity) could easily be visualised (Figure 7). Additionally, many subcellular features of sporozoites — the apical polar ring, rhoptries, parasite plasma membrane, and basal complex — could be easily distinguished just by using protein-density dyes (NHS ester) (Figure 7). These same features could not be determined in unexpanded sporozoites using the same fluorescent dyes. We learned that reimaging the same salivary gland before and after expansion did not provide a clear anatomical comparison as it did for midguts. Although we successfully imaged the same salivary gland before and after expansion (Figure 3a), this method presented challenges. Each lobe of the salivary gland had independent mobility from its original imaging to expansions. Therefore, it was challenging to correlate the anatomical structures between the initial and expanded states of the salivary glands (Figure 5).

**Figure 6:**
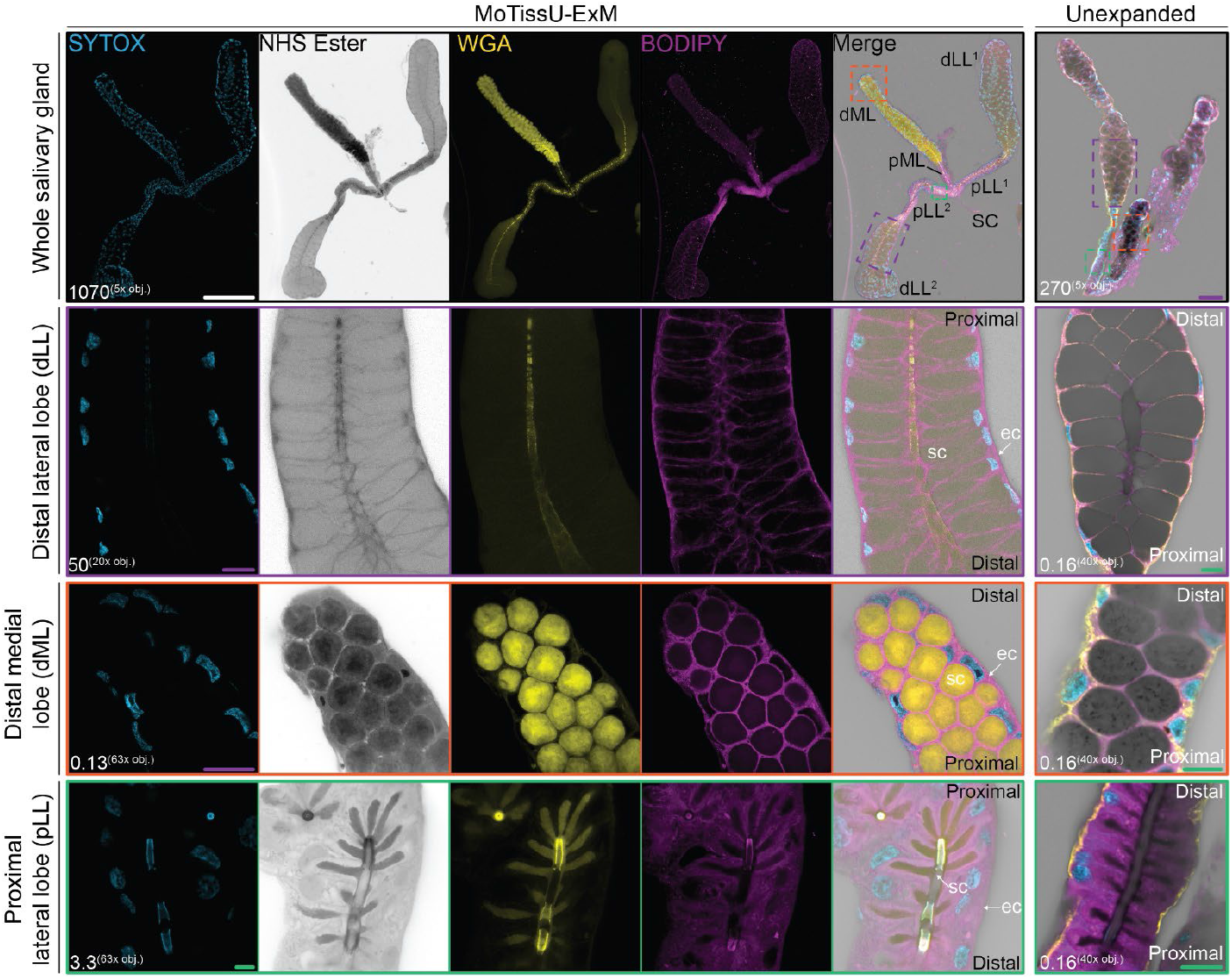
U-ExM preserves salivary gland anatomical structure and ultrastructure. Mosquito salivary glands were prepared by U-ExM, or left unexpanded, stained with SYTOX (cyan, DNA), NHS ester Alexa Fluor 405 (greyscale, protein density), wheat germ agglutinin Texas red (WGA (yellow, chitin & glycans), and BODIPY FL-Ceramide (magenta, lipids), and imaged by Airyscan Microscopy. In U-ExM salivary glands, all anatomical features were preserved including secretory cavity, and both the distal (d) and proximal (p) regions of the medial lobe (ML) and lateral lobes (LL). Zoomed in regions of a distal lateral lobe (dLL^2^, magenta), the distal medial lobe (dML, orange), and a proximal lateral lobe (pLL^2^, green) show their ultrastructural preservation and the clear distinction between the epithelial cell cytoplasm (ec) and secretory cavity (sc). Proximal and distal are indicated on zoomed regions to show their orientation. Note that the unexpanded salivary gland is not the same salivary gland prepared by U-ExM. Number in the bottom corner of each image indicates z-projection depth in µm. obj.= objective lens. Scale bars: = white 500 µm, magenta = 50 µm, green = 10 µm.

**Figure 7:**
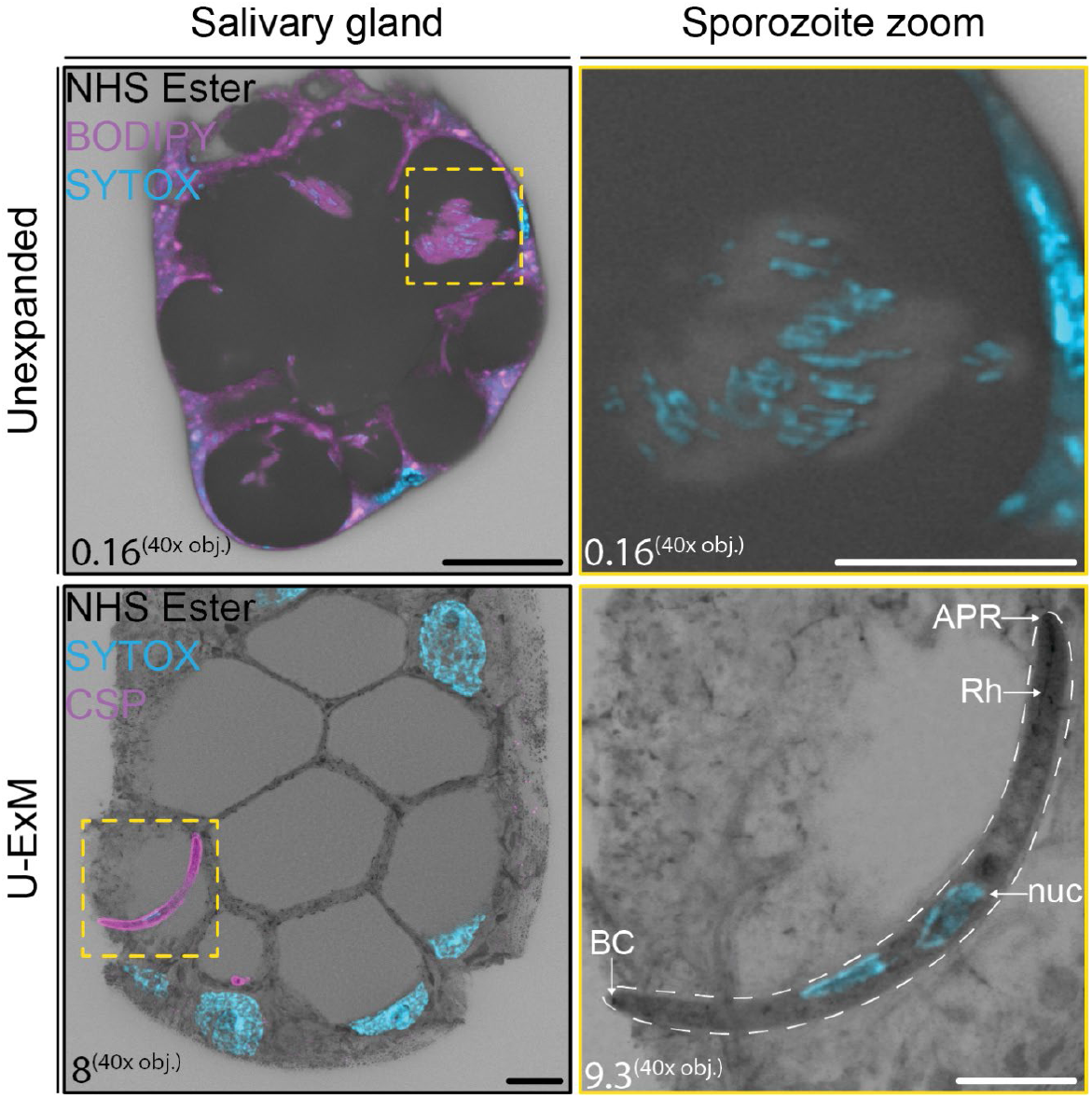
Comparison of sporozoites in unexpanded and U-ExM salivary glands. Mosquito salivary glands infected with *P. berghei* were fixed, and either imaged unexpanded or prepared by U-ExM. Unexpanded salivary glands were stained with SYTOX (cyan, DNA) and NHS ester Alexa Fluor 405 (greyscale, protein density), and BODIPY-FL-Ceramide (magenta, lipids). U-ExM salivary glands were stained with STYOX, NHS ester and anti-circumsporozoite protein (CSP) antibodies (magenta, sporozoite surface). Depicted is a section of the infected salivary gland, along with a zoom of sporozoites inside the salivary gland (yellow). In salivary glands prepared by U-ExM, the nucleus (nuc), parasite plasma membrane (white dashed line), basal complex (BC), rhoptries (Rh), and apical polar ring (APR) can all be visualised. Number in the bottom corner of each image indicates z-projection depth in µm. obj.= objective lens. Scale bars: black = 20 µm, white = 10 µm.

### Addressing challenges of sample depth

For the implementation of this protocol, the most significant technical hurdle we faced was imaging expanded tissues, especially salivary glands, due to the depth of the expanded sample combined with the shallow limit of the working distance of an oil immersion objective. This means that typically, only a very small portion of the expanded mosquito tissue is accessible using a standard high-resolution imaging setup. A typical high-resolution confocal microscope setup would include a high numerical aperture (NA) oil-immersion objective (such as the 63x, 1.4 NA objective used in this study), with a working distance of < 200 µm. Following expansion, the distance from the start of the gel to the basal side of the tissue is always significantly greater than 200 µm (typically 500 – 1000 µm). This effect is amplified in salivary glands, which are frequently suspended within the gel rather than lying flat on one surface.

To mitigate this limitation for mosquito midguts, we first tried longitudinally ‘opening’ the midgut following dissection. We reasoned that opened midguts would lay flat within the gel, with both sides imageable on a high-resolution imaging setup. Longitudinally opened midguts were compatible with U-ExM (Figure 8) and, in some instances dramatically increased the area of the expanded midgut accessible using a 63x oil-immersion objective lens. In many instances, however, the edges of the longitudinally opened midgut curled and made even less of the tissue accessible using a 63x oil-immersion objective lens.

**Figure 8:**
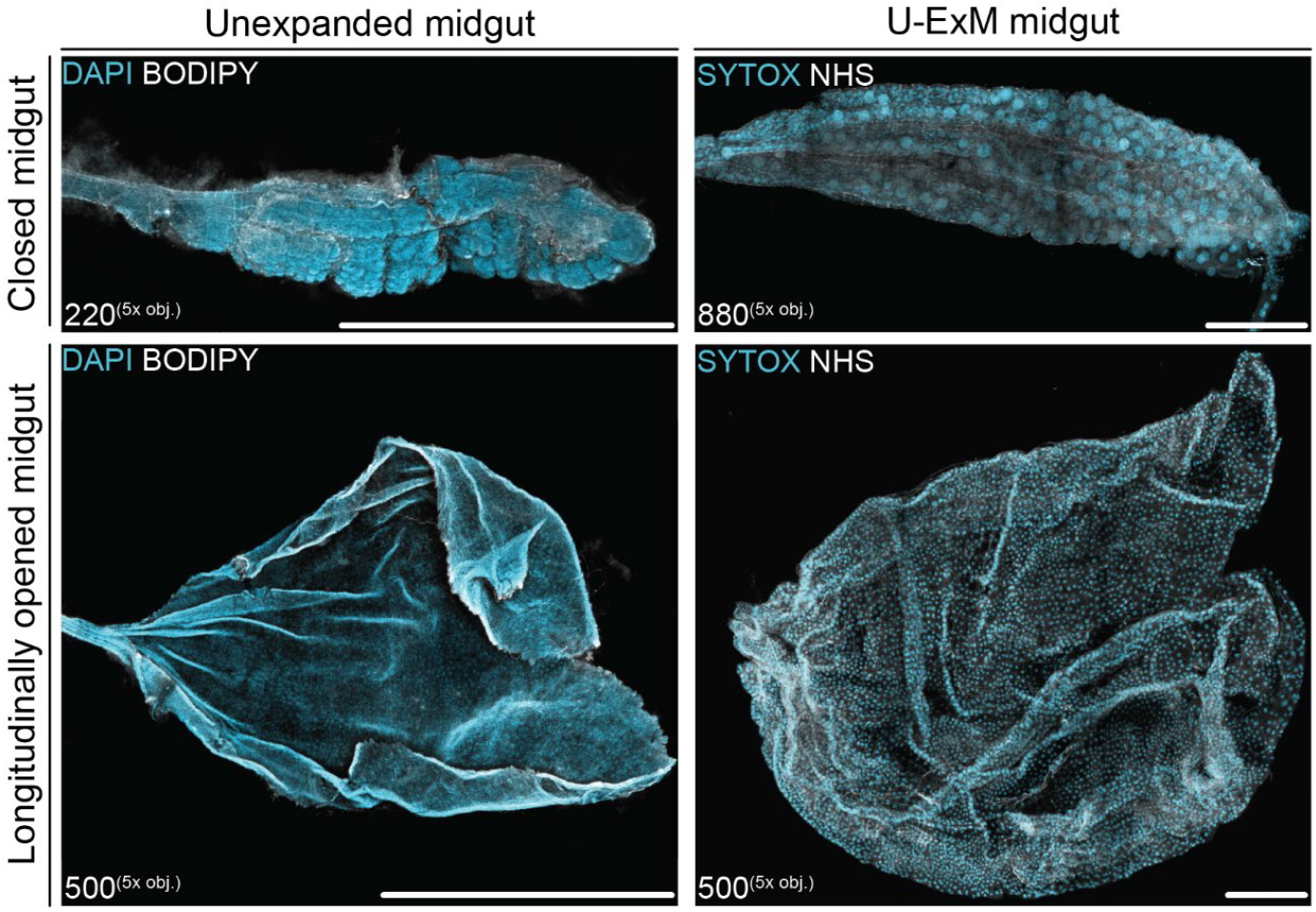
Longitudinally opened midguts prepared using MoTissU-ExM. During mosquito tissue dissection, midguts were either removed intact (closed) or ‘opened’ longitudinally. Opened midguts were prepared for U-ExM identically to closed midguts. Unexpanded midguts were stained with DAPI (cyan, DNA) and BODIPY-TR-Ceramide (white, lipids), while U-ExM midguts were stained with SYTOX (cyan, DNA) and NHS Ester Alex Fluor 405 (white, protein density). Number in the bottom corner of each image indicates z-projection depth in µm. obj.= objective lens. Scale bar = 1000 µm.

While we could frequently observe *P. berghei* oocysts within an expanded midgut, using a 63x oil-immersion objective lens, we could never image the entirety of the oocyst depth as it would exceed the objective working distance (Figure 9a). To overcome this, we utilised a 40x water-immersion (1.2 NA) objective lens with a ∼500 µm working distance. Using this 40x objective, we were able to measure the depth of an oocyst (Figure 9b) that we could not measure using the 63x oil-immersion objective. Additionally, despite the 40x water-immersion objective having a lower NA, the image was markedly brighter and clearer than using the 1.4 NA 63x objective (Figure 9). This is to be expected as the NA of an objective assumes refractive index matching between immersion and the sample. As the gel is mostly water, this means the effective NA of the 63x oil-immersion is <1.4, and in this instance likely marginally lower than the effective NA of the 40x water-immersion objective lens.

**Figure 9:**
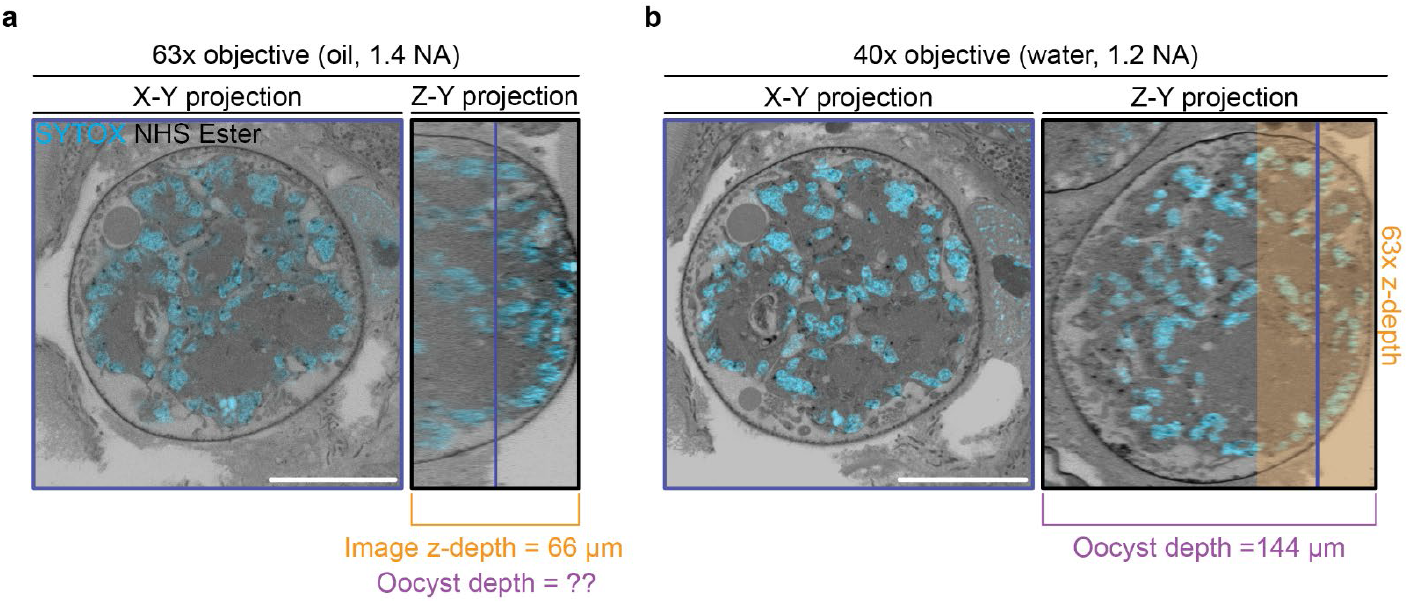
Comparison of oil and water immersion objectives for imaging expanded samples. A *P. berghei* infected mosquito midgut was prepared by U-ExM, stained with SYTOX (cyan, DNA) and NHS ester Alexa Fluor 405 (greyscale, protein density), and imaged by Airyscan Microscopy. **(a)** Image of an oocyst using a 63x oil-immersion objective showing a single X-Y slice and Z-Y projection (3D image rotated 90° to the right). Using this imaging setup, the image z-depth reached 66 µm before exceeding the working distance of the objective. Oocyst depth could not be measured, as it exceeded the image z-depth. **(b)** The same oocyst imaged using a 40x water-immersion objective with a longer working distance. Using this imaging setup, the oocyst depth was measured at 144 µm. The blue line indicates the z-axis position of the z-slice depicted in the X-Y projections. Scale bar = 50 µm.

## Discussion

Here, we developed the first protocol for *in situ* visualisation of mosquito tissues and the malaria parasites within them using ultrastructure expansion microscopy (MoTissU-ExM). Although this protocol was primarily developed for the purpose of imaging malaria parasites along with their mosquito hosts *in situ*, this protocol holds broader applications for mosquito biology research. It could be used to study mosquito tissues in isolation or to study their interaction with mosquito-borne pathogens, such as *Flaviviridae* and *Aedes* mosquitoes, or filarial worms and *Culex* mosquitoes. More broadly, this protocol could be applied to any vector whose midgut and salivary gland(s) can be easily dissected, including sandflies infected with *Leishmania*, blackflies with *Onchocerca*, tsetse flies and triatomines with trypanosomes, or ticks with any of the various bacteria, viruses, and parasites they transmit.

### Use of longitudinally opened midguts

We showed that longitudinally opened midguts can be prepared using the MoTissU-ExM protocol. When longitudinally opened, approximately half of prepared midguts would be oriented in a gel in way that dramatically increased the midgut surface area accessible for imaging. The other half, however, would be oriented with the ‘opened’ side facing towards the cover glass (towards the objective). When in this orientation, almost none of the midgut would be accessible for imaging at high magnification. For example, the unexpanded opened midgut in Figure 8 required a z-depth more than twice as deep as the closed midgut to capture the entire tissue. However, even considering this limitation there may be some instances where the longitudinal opening of midguts would be favourable for MoTissU-ExM, *i*.*e*. when burden of parasites per midgut is high, or the number of midguts is not a limiting factor. In this study for example, we exclusively used *P. berghei*, which typically reaches high midgut oocyst burdens well over 100 per midgut^25^. In cases where the oocyst burden is substantially lower, such as for *P. falciparum*^*26*^ or a mutant/drug treated parasite, but the numbers of midguts is not a limiting resource, longitudinally opening midguts may be favourable. By contrast, if the numbers of midguts available for imaging is low, it would not be favourable to longitudinally open these midguts, as approximately 50% of them will not contain imageable oocysts.

### Common challenges of imaging MoTissU-ExM samples

The primary challenge in imaging MoTissU-ExM samples lies in the depth of the sample, which poses difficulties for high-resolution imaging. Using a conventional high-resolution objective lens with a high-numerical aperture (1.4 NA), we consistently encountered limitations to image an entire oocyst without exceeding the working distance of the lens. Additionally, we were often unable to access any of the expanded tissue using this imaging setup. Hence, it is advisable to always image samples using long working distance objective lenses. In our opinion, any compromise in resolution due to the use of a lower NA, longer working distance lens, is offset by improved sample accessibility. Further, long working distance objectives, such as the 40x objective lens used in this study, are frequently water-immersion objectives. Given that the expanded gel is almost entirely water, using a water-immersion objective lens helps minimise spherical aberration compared to oil-immersion objectives (visible in Figure 9). Alternatively, MoTissU-ExM samples could be imaged using methods where sample depth is not a significant concern, such as using LightSheet fluorescence microscopy^27^.

It has previously been shown that the presence of chitin in tissues can limit expansion, which can be overcome by treating samples with chitinase^22^. In this protocol, we did not treat tissues with chitinase and despite this no significant aberrations in expansion were noticed either at the tissue or cell level. This is likely because the midgut and salivary gland have relatively low amounts of chitin by comparison to the exoskeleton or wing tissue, for example. Within the salivary glands, however, the secretory duct is thought to be highly chitinous^28^ and indeed the secretory duct was highly fluorescent using wheat germ agglutinin (WGA), which binds chitin and sialic acid (Figure 6). To the best of our understanding, chitin should not expand and should not be anchored to the gel using U-ExM so it is unclear why WGA fluorescence is strong in regions thought to be chitinous. One possibility is that in these expanded samples, WGA is not binding chitin and instead the fluorescence of WGA corresponds to sialylated proteins, and that these happen to be similar to the distribution of chitin. It is noteworthy that while the secretory duct is also highly protein dense, the WGA signal is not identical to protein density (Figure 6), which is a known trait of fluorescent dyes with high background^14^. Currently, the mechanism by which molecules are crosslinked to hydrogels using formaldehyde and acrylamide is not entirely understood^29-31^ and so it is unclear whether WGA fluorescence corresponds to chitin, sialylated proteins, or something else.

In the development of MoTissU-ExM, we typically used highly infected midguts and salivary glands to facilitate the detection of parasites within the tissue. When parasite burdens are high, oocysts can easily be found in midguts and sporozoites in salivary glands using either DNA or protein density dyes. In cases with low parasite burden, we recommend the use of a parasite-specific marker (such as an anti-capsule antibody or anti-circumsporozoite protein antibody for example), which will allow unambiguous identification of parasites within tissues. In expanded tissues, fluorescent parasites will be visible even at low-magnification so large areas of the tissue can be scanned for parasite-specific fluorescence at low magnification before switching to a high-magnification objective lens for imaging.

### Adaptations of U-ExM and iterative U-ExM methods

Our development of MoTissU-ExM mostly focussed on how to dissect mosquito tissues and reliably get them into hydrogels. Considering this, the protocol should be adaptable to the many other forms of expansion microscopy, or experiments multiplexed with expansion microscopy. For example, this protocol could simply be adapted for ten-fold robust expansion microscopy (TREx)^32^, MAGNIFY^33^, iterativeU-ExM (iUExM)^34^, or ExFISH^35^. One caveat to this is that issues of sample depth would be considerably worse for expansion microscopy protocols that result in greater than 4-fold one-dimensional expansion. Therefore, if TREx, MAGNIFY, or iUExM were applied to mosquito tissues, for example, it is likely that LightSheet fluorescence microscopy would need to be used to access the sample. One possible alternative to overcoming the issue of sample depth, would be to make transverse sections of a gel and then image the sections. To the best of our knowledge, however, this has not been previously published.

## Methods

### Image acquisition

All images in this study were acquired using an LSM900 AxioObserver with Airyscan 2 (Zeiss, Oberkochen, Germany). Images were acquired on either a 5x Fluar (air, 0.25 NA), 20x Plan-apochromat (air, 0.8 NA), 40x C-apochromat autocorr M27 (water, 1.2 NA), or 63x Plan-apochromat (oil, 1.4 NA). The objective used for each image in this study is indicated in the bottom left corner of the image.

### Image processing and presentation

All images in this study underwent Airyscan processing using ‘moderate’ filter strength. Images acquired using the 5x and 20x objectives underwent 2D Airyscan processing, while images acquired using either the 40x or 63x objectives underwent 3D Airyscan processing. For images that contain NHS Ester, the gamma value of this channel was set to 0.45, rather than 1 for greater discernment of subcellular structures; as has previously been described^14^.

All images in this study were prepared and processed in, then exported from ZEN Blue Version 3.5 (Zeiss, Oberkochen, Germany). Images in Figures 2a, 3a, 5b, 6, 7, & 10 were rotated and/or cropped from a larger original image to aid comparison.

### Unexpanded tissue imaging

Unexpanded midguts or salivary glands were dissected and fixed as described in the point-by-point protocol. Fixed tissues were then washed in PBS to remove the fixative and stained with Hoechst, SYTOX, NHS Ester Alexa Fluor 405, or BODIPY-FL-Ceramide diluted in PBS for 1 hour at room temperature. Stained tissues were then transferred to Poly-D-Lysine coated 35 mm imaging dishes with 22 mm #1.5 cover glass bottoms, covered with ProLong Glass and imaged using Airyscan Microscopy.

### Imaging of tissues before and after U-ExM

For the salivary glands and midguts that were imaged both before and after U-ExM, the tissues were initially prepared as described for unexpanded tissue imaging except without Poly-D-lysine coating the imaging dishes. These tissues were then imaged immediately after application of ProLong Glass. Once imaging was complete, imaging dishes were washed multiple times with PBS until the tissues detached from the imaging dish. Detached tissues were further washed in PBS overnight. Once washed, tissues were transferred to anchoring solution and prepared for U-ExM as described in the point-by-point protocol. During imaging, these tissues were intentionally oriented in a manner similar to the unexpanded image to aid comparison.

### Measurement of nucleus diameter

To measure nucleus diameter, the slice at which each nucleus had its maximum diameter was found and this diameter was measured manually in 2D using the ‘Profile’ function of ZEN Blue. Only one diameter measurement was made per nucleus. Measurement histograms were imprinted on top of measured nuclei to prevent repeated measurements of the same nucleus.

### Statistical analysis

All graphs presented in this study were generated using GraphPad PRISM (version 10). All error bars represent standard deviation. On both graphs, large datapoints represent mean values of nucleus diameter from a single tissue, while small datapoints represent diameter measurements of individual nuclei.

1D-expansion factor was estimated by dividing the mean expanded nucleus diameter (31.62 µm for midguts, 42.63 µm for salivary glands) by the mean unexpanded nucleus diameter (7.25 µm for midguts, 10.17 µm for salivary glands).

### Stains and antibodies

A list of stains and antibodies used in this study, along with their working concentrations, source and step of application is listed in Tables 1 and 2.

**Table 1:** Summary of all antibodies used in this study.

| Antibody (Ab) | Ab species | Ab source | Ab concentration | Step of protocol applied | Reference/cat no. |
| --- | --- | --- | --- | --- | --- |
| Anti-*Pb*circumsporozoite protein (3D11) | Mouse | BEI Resources | 1: 1000 | Primary Ab | MRA-100A<sup>36</sup> |
| Anti-mouse IgG Alexa Fluor 555 (H+L) | Goat | Invitrogen | 1:500 (2mg/ml stock) | Second Ab | A21422 |
| Anti-mouse IgG Alexa Fluor 488 Superclonal (H+L) | Goat | Invitrogen | 1:500 (1mg/ml) | Second Ab | A28175 |

**Table 2:** Summary of all fluorescent dyes and stains used in this study. *Indicates that dye was also used on unexpanded samples.

| Stain/dye | Source | Concentration | Step of protocol applied | Cat no. |
| --- | --- | --- | --- | --- |
| NHS Ester Alexa Fluor 405 | ThermoFisher | 1:250 (2 mg/mL stock in DMSO) | Secondary Ab* | A30000 |
| BODIPY-TR-Ceramide | ThermoFisher | 1:500 (1 mM stock in DMSO) | Post expansion | D7540 |
| BODIPY-FL-C<sub>5</sub>-Ceramide | ThermoFisher | 1:500 (1 mM stock in DMSO) | Post expansion* | D3521 |
| SYTOX Deep Red | ThermoFisher | 1:1000 (1 mM stock in DMSO) | Secondary Ab* | S11381 |
| Wheat Germ Agglutinin Texas Red-X | ThermoFisher | 1:250 (1 mg/mL stock in PBS) | Post expansion | W21405 |
| Hoechst 33342 | ThermoFisher | 1:1000 (10 mg/mL stock in water) | Unexpanded only | C10329G |

## Point-by-point protocol for <u>Mo</u>squito <u>Tiss</u>ue <u>U-ExM</u> (MoTissU-ExM)

### Background

This protocol is ultimately based on the first U-ExM protocol^4^, and the corresponding adaptations made to that protocol for imaging of malaria parasites^6^, paraformaldehyde-fixed malaria parasites^37^, along with optimisations for anchoring time^21^ and data presentation^14^.

### Reagents & chemicals

- Deionised, nuclease-free water
- Poly-D-Lysine 0.1mg/mL solution (Gibco cat no: A3890401)
- Ammonium persulfate powder (APS, ThermoFisher cat no: 17874)
- Tetramethylethylenediamine (TEMED, ThermoFisher cat no: 17919)
- Formaldehyde 36.5-38% solution (FA, Sigma cat no: 8775)
- Acrylamide 40% solution (AA, Sigma cat no: A4058)
- N,N’-methylenebisacrylamide 2% (BIS, Sigma cat no: M1533)
- Sodium acrylate >97% powder (SA, Sigma cat no: 408220)
- Propyl gallate 98% powder (ThermoFisher cat no: 131581000)
- Tris (hydroxymethyl) aminomethane (TRIS Base, RPI cat no: T60040-250.0)
- Sodium dodecylsulfate micro-pellets (SDS, RPI cat no: L22040-500.0)
- Sodium chloride (NaCl, RPI cat no: S23020-1000.0)
- Glycerol (Fisher cat no: BP229-4)
- 10x PBS (Sigma cat no: 806552)
- Bovine serum albumin
- TWEEN-20 (Sigma cat no: P1379)

### Solutions

- <u>Sodium acrylate solution</u> (38% wt/wt in Milli-Q water)
  ∘ WARNING: sodium acrylate solution should be prepared and used in the fume hood due to the toxicity of powdered sodium acrylate.
- <u>TEMED solution</u> (10% in Milli-Q water)
  ∘ WARNING: TEMED Solution should be prepared and used in the fume hood due to the toxicity of concentrated TEMED.
  ∘ Aliquot and store at -20 °C
- <u>APS solution</u> (10% in Milli-Q water)
  ∘ Aliquot and store at -20 °C
- <u>Monomer solution</u> (sodium acrylate 19% wt/wt, acrylamide 10% v/v, BIS 0.1% v/v, in PBS)
  ∘ WARNING: Monomer solution should be prepared and used in the fume hood.
  ∘ Monomer solution MUST be prepared at least 24 hours in advance and can be stored at - 20 °C for up to 2 weeks.
  ∘ We typically make 900 µL of monomer solution as follows, and store as 10 x 90 µL aliquots (good for making 20 gels)
    ▪ 500 µL 38 % wt/wt sodium acrylate solution
    ▪ 250 µL acrylamide
    ▪ 50 µL BIS
    ▪ 100 µL 10x PBS
- SDS stock solution (350 mM in Milli-Q water)
  ∘ WARNING: SDS stock solution should be prepared in the fume hood due to the toxicity of powdered SDS.
- NaCl stock solution (5 M)
- <u>Denaturation buffer</u> (200 mM SDS, 200 mM NaCl, 50 mM Tris, pH 9, in water)
  ∘ WARNING: Gloves should always be worn when handling denaturation buffer as it is highly irritating to skin.
  ∘ We typically make 500 mL of denaturation using the following recipe:
    ▪ 3 g TRIS base
    ▪ 285.7 mL SDS stock solution
    ▪ 20 mL NaCl stock solution
    ▪ Adjust to pH 9 with HCl
    ▪ Fill to 500 mL with Milli-Q water
  ∘ Due to the high concentration of SDS, the denaturation buffer often falls out of solution, but this can be easily reversed by gently heating.
- <u>Anchoring solution</u> (1.4 % v/v formaldehyde, 2 % v/v acrylamide in PBS)
  ∘ WARNING: Anchoring solution should be prepared and used in the fume hood.
  ∘ Should be made fresh for each experiment
- Propyl gallate solution (0.2 % wt/v propyl gallate in Milli-Q water)
- <u>Freezing solution</u> (50% v/v glycerol in Milli-Q water)
- <u>Blocking solution</u> (3% w/v bovine serum albumin in PBS)
- <u>Wash buffer</u> (0.5% v/v TWEEN-20 in PBS)

### Materials

- Smart plastic razors (Sigma cat no: Z740503)
- Thin paintbrush (#2 to #4 brush width)
- 1.5 mL tube cap locks (Fisher cat no: NC9679153) or locking 1.5 mL tubes (ThermoFisher cat no: 3456)
- 12 mm round coverslips (Fisher cat no: NC1129240)
- 35 mm Cellvis #1.5 glass bottomed dishes (Fisher cat no: NC0409658)
- Parafilm (Sigma cat no: P7793)
- 37 °C incubator
- Dry heat block
- Orbital shaker
- Vortex
- Humid chamber
- Dissecting forceps (Fisher cat no: NC9889584)

### Mosquito rearing and dissection

1. *An. gambiae* (Keele strain)^38^ mosquitoes were maintained at 27 °C and 80% relative humidity, following a 14-hour/10-hour light/dark cycle under standard laboratory conditions at the National Institutes of Health.

### Mosquito Infections

1. Infections were performed using a *Plasmodium berghei* ANKA strain genetically modified to express a green fluorescent protein (GFP) driven by the elongation factor 1A (ef1α) promoter^39 40^) in the background of either *ron11*^KO^ or *ron11*^ctrl^parasites.
  i. Allow female mosquitoes, aged 4-5 days, to feed on *P. berghei*-infected mice with an exflagellation rate of 3-4 exflagellants per 40x microscopic field.
  ii. Perform oviposition: after three to four days at 19°C, the mosquitoes were allowed to lay eggs. NOTE: This step is critical if you want to perform an additional feeding.

### Sample preparation & fixation

#### Midguts

1. Dissect mosquito midguts in phosphate-buffered saline (PBS, 130 mM NaCl, 7 mM Na_2_HPO_4_, 3 mM NaH_2_PO_4_·H2O pH 7.2) at room temperature.
2. Immerse the midguts straight in 4% paraformaldehyde in PBS after dissection.

### Longitudinally opened midguts

1. To longitudinally open midguts, allow mosquitoes to feed on freshly prepared 10% BSA in 0.15 M Sodium Chloride mixed with 10 mM sodium bicarbonate, pH 7.2^41 42^.
  a. The pH **must** be adjusted right before feeding.
2. Fix the midguts for 30 s with 4% paraformaldehyde (PFA) to preserve the midgut structure.
3. Open the midguts longitudinally using a pair of number 5 dissecting forceps.
4. After opening, fix the midguts for 1 hour with 4% PFA in PBS.

#### Salivary glands

1. Dissect the salivary glands in PBS at room temperature.
2. Immediately fixed with 4% PFA in PBS.

### Anchoring

1. After fixation, remove as much fixative as possible form the 1.5 mL tube containing fixed mosquito tissues; being careful not to remove any of the fixed sample.
  i. It is best to do this with a P200 or smaller pipette, and ideally you will leave < 100 µL of fixative in your tube.
  ii. To avoid mosquito tissues adhering to the inside of the tip if accidentally pipetted, you can coat the inside of your tip with <u>blocking solution</u> before removing the fixative.
2. Fill tube with <u>anchoring solution</u> (1.2% AA, 2% FA) and incubate overnight at 37 °C.

### Gelation

1. Prepare <u>monomer solution</u>
  i. IMPORTANT: <u>Monomer solution</u> must be prepared at least 24 hours before gelation.
2. Coat 12 mm round coverslips with poly-D-lysine for 1 hour at 37 °C before washing twice with Milli-Q water.
3. Thaw aliquots of <u>APS solution</u> and <u>TEMED solution</u> on ice for 30 minutes
4. Prepare a humidity chamber that contains one square of parafilm for each gel you will make and place at -20 °C for 20 minutes.
  i. We typically use 30 mL Petri dishes containing a single wet Kimwipe as a humidity chamber, these can hold four squares of parafilm.
5. Place humidity chamber on ice, along with one aliquot of <u>Monomer solution</u> for every two gels you will make.
6. Place a poly-D-lysine coated coverslip onto the parafilm square in the humidity chamber.
7. Remove anchoring solution from fixed mosquito tissues, concentrating the sample in as little anchoring solution as possible.
  i. We typically aim to transfer ∼30-50 µL of solution containing the samples to each coverslip.
8. Cut a P200 pipette tip with scissors to widen its opening and coat the inside of the tip with <u>Blocking solution</u> to prevent tissues from sticking to the tip.
9. Resuspend ∼5 salivary glands or midguts in the remaining anchoring solution and transfer to the poly-D-lysine coated coverslip.
  i. The number of salivary glands or midguts per gel can be varied depending on the experiment (see optimisation section), but we find that this number of tissues is best to ensure no two tissues are overlapping in the gel.
10. Using a P10 pipette, gently remove the remaining anchoring solution from the coverslip, being careful not to remove any of the tissues.
  i. While it is best to remove as much anchoring solution as possible, you do not want your tissues to dry out.
11. Add 5 µL of <u>TEMED solution</u> and 5 µL of <u>APS solution</u> to an aliquot of <u>Monomer solution</u>, very briefly vortex (1-2 seconds) and pipette 35 µL onto the parafilm for each gel.
  i. Two gels can be made from each aliquot of <u>Monomer solution</u>.
12. Using forceps, pick up the coverslip containing the mosquito tissues, invert it (so the tissues are facing down), and gently place on top of the bubble of monomer solution.
13. Leave samples on ice for 10 minutes for gel to start solidifying.
14. Transfer humidity chamber to 37 °C for 30 minutes to allow gel to polymerise.

### Denaturation

1. Set a heat block to 95 °C.
2. Fill the wells of a 6-well plate with 2 mL of <u>denaturation buffer</u>.
3. Transfer gel and coverslip from the parafilm to a well containing denaturation buffer and place on an orbital shaker for 15 minutes at room temperature to allow the gel to separate from the coverslip.
  i. Gel will detach more easily if it is placed coverslip-side down.
4. Once gel has detached from coverslip, remove from the 6-well plate and place it on a flat surface. At this point, mosquito tissues should be visible in the gel.
5. Using a smart plastic razor, cut around each of the tissues so they can be processed individually.
  i. NOTE: This step is not necessary, however, it can significantly streamline the staining and imaging process (see Optimisation section).
  ii. Alternatively, the tissues can be cut from the gel as round pieces using a circle punch cutter or the wide end of a pipette tip.
6. Fill a 1.5 mL tube with <u>denaturation buffer</u> and place each of the gel pieces containing a tissue into a separate tube.
  i. NOTE: These small gel pieces can be difficult to manipulate, we find it easiest to handle them using a paintbrush (size #2 - #4).
  ii. Instead of 1.5 mL Eppendorf tubes, smaller volume screw-cap or locking lid tubes can also be used.
7. Place a safety cap onto each 1.5 mL tube and denature the samples in the 95 °C heat block for 90 minutes.

### First expansion

1. Prepare 6-well plates where for each tissue a well contains 2 mL of MilliQ water.
2. Following denaturation, remove denaturation buffer from the tube and transfer each tissue to a well of the 6-well plate.
  i. CAUTION: Tubes will be very hot coming out of the heat block, remove from heat block with heat-resistant gloves and allow to cool before handling.
  ii. When removing the denaturation buffer from the tube, it is best to use a P200 pipette, as the gel can easily be sucked into a P1000 pipette tip.
3. Place 6-well plate onto an orbital shaker and allow gels to expand for 30 minutes at room temperature.
4. After 30 minutes, remove the water and perform two more 30-minute water washes to fully expand the gel.
  i. NOTE: At this point the gels can be cryopreserved (see Storage, transportation & cryopreservation of samples section for details).

### Staining

1. Shrink fully expanded gels down by performing two 15-minute washes in 2 mL of 1 x PBS.
2. Transfer shrunken gels to the wells to a 12-well or 24-well plate.
  i. This step is not strictly necessary but will significantly reduce the amount of antibodies and stains used per gel. For 12-well plates, block or wash using 1 mL of solution and stain using 500 µL of solution. For 24-well plates, block or was using 500 µL of solution and stain using 250 µL of solution.
3. Place samples in <u>Blocking solution</u> for 30 minutes at room temperature on an orbital shaker.
4. Prepare primary antibodies in <u>Blocking solution</u> and add to gels overnight at room temperature.
  i. For antibodies that have not been used in U-ExM experiments before, we tend to start by using 2x the concentration that would be used for conventional immunofluorescence microscopy. For a list of working concentrations of common antibodies and dyes see the relevant section below.
5. Following primary antibody staining, wash the gels three times in <u>Wash buffer</u> for 10-minutes each.
6. Prepare secondary antibodies in 1 x PBS and add to gels for 2.5 hours at room temperature.
  i. IMPORTANT: Secondary antibodies can be prepared in blocking solution instead of PBS, but this will dramatically reduce the fluorescence of protein dyes like NHS Ester.
  ii. NOTE: Fluorescent dyes that are not conjugated to antibodies, like DAPI or NHS ester are typically included with this secondary antibody incubation.
7. Following secondary antibody staining, wash the gels three times in <u>Wash buffer</u> for 10-minutes each.
8. After washing, move gels back the wells of a 6-well plate and re-expand by washing three times for 30-minutes in Milli-Q water.

### BODIPY and WGA staining

1. For gels that will be stained BODIPY ceramide, or wheat germ agglutinin (WGA), remove the water from the gel and place in the dye diluted in 1mL of 0.2 % w/v propyl gallate per well.
  i. Propyl gallate is not necessary here, and the dye can instead be diluted in Milli-Q water, but propyl gallate has a modest antifade effect and can improve fluorophore brightness and longevity.
  ii. Lipid dyes are extremely sensitive to washing on expanded samples and will lose their fluorescence with any significant washing.

### Sample mounting

Note: This protocol is optimised for inverted microscopes, however a guide for preparing U-ExM gels to image using an upright microscopy can be found here^43^.

1. For each gel, prepare a glass-bottomed imaging dish by coating it with ∼500 µL Poly-D-lysine solution and incubating for 30 minutes at 37 °C
2. Remove gel from 6-well plate and place on a flat surface.
  i. In their fully expanded state, gels are best handled using a smart plastic razor or similar.
3. Determine the sidedness of the gel and place the gel into the imaging dish with the ‘coverslip side’ facing down.
  i. During gelation the gel has one side that forms in contact with the coverslip (where the tissue is), and a side that forms in contact with the parafilm. When closely inspected the side that contacted coverslip typically appears perfectly smooth, while the side that contacted the parafilm typically has a slightly ruffled appearance form the pattern on the parafilm.
  ii. To minimise the sample drifting, it can also be helpful to gently press the gel onto the imaging dish to ensure the entire gel contacts the glass surface and to minimise air bubbles.

### Storage, transportation & cryopreservation of samples

Following either the first expansion (unstained) or second expansion (stained), gels can cryopreserved as follows^34^:

1. Prepare <u>Freezing solution</u> (50% v/v glycerol in MillIQ water).
  i. Approximately 3 mL freezing solution will be needed per gel.
2. In the wells of a 12-well plate, wash gels three times for 30 minutes in 1 mL <u>Freezing solution</u> on an orbital shaker at room temperature.
3. Place gels at -20 °C for long term storage.
  i. Gels that were cryopreserved unstained are stable at -20 °C for multiple years. Gels that were cryopreserved following staining start to become noticeably less fluorescent after 6-12 months of storage.

Development of this technique required shipment of mosquito tissues. To do this, fixed tissues were simply placed in 1x PBS after fixation and stored in locking 1.5 mL tubes. These samples were then secured and shipped on ice. From this process, we noticed no damage to the transported tissues. Fixed tissues were stable at 4 °C for a few weeks before anchoring and gelation. In addition, we successfully transported cryopreserved gels on ice in 12-well plates that had been sealed closed with Parafilm.

### Optimisation

#### Tissues number per gel and gel processing

In the development of this protocol, we were typically working with highly-infected tissues where each tissue had dozens to hundreds of imageable parasites. Because of this, only relatively few tissues were required to get many images, leading us to process and stain each tissue individually. For experiments using either lowly-infected tissues, or looking addressing tissue-level biological questions, it may be desirable increase the number of tissues per gel or process tissues in batches rather than individually. In this protocol, we suggest ∼5 tissues per gel and to cut them out and stain them individually. If a greater number of tissues is required, or tissue overlap is not a particular concern however, the number of tissues per gel can comfortably be increased. Further, rather than being processed individually, a whole gel could be stained to look at many tissues at a time, or pieces of gel could be cut containing more than one tissue and stained as a group.

#### Increasing throughput

Placing infected tissues into gels then cutting them out individually and processing them can be very time consuming. We found the best way to increase our throughput in this regard was to place all tissues from a given experiment into gels at once and cryopreserve them unstained. That way, we could image all the replicates from a given experiment in one batch.

In this study, we recommended to place ∼5 tissues per gel. We found it helpful to streamline this step during tissue dissection by fixing dissected tissues in multiple 1.5 mL tubes each contain 5 tissues. This way, each gel corresponds to a particular 1.5 mL of dissected tissues, rather than a portion of the tissues coming from a tube that contained many more tissues.

### Imaging tips

- From the illumination of a microscope lamp, especially the channel corresponding to NHS ester, intact tissues can easily be seen by eye. We found it useful to utilise this to orient the tissue, longitudinally for example, to streamline whole-tissue imaging (Figure 10).
- When imaging whole tissues, we find it best to first image the entire intact tissue on a low magnification objective lens (such as 5x) before moving onto a high magnification objective lens (such as 63x) to image individual oocysts or sporozoites. Doing this will allow for pairing of ultrastructure-level imaging data with the anatomical context of the parasite.
- High magnification and high numerical aperture (NA) objective lenses typically have a free working distance that is poorly compatible with expanded tissues. For example, using a 63x (1.4 NA) oil immersion objective lens, we were never able to image an entire oocyst without exceeding the free working distance of the objective lens (Figure 9). We had far greater success imaging our samples with long working distance water immersion objectives. The trade-off in NA (and therefore resolution) was marginal considering how much more of the sample could be accessed.
- Dyes and stains that are not specific to individual proteins, like BODIPY ceramide, WGA, or NHS ester can be very useful for orientation and context purposes. We would recommend always using at least one of these kind of stains.
- As expanded gels are mostly water, imaging using oil-immersion objective lenses can lead to spherical aberration. Considering this, it is best to image using water-immersion objective lenses if possible.
- Finding sporozoites, even within a highly infected salivary gland, can be very challenging without a sporozoite-specific marker. We typically used antibodies against the circumsporozoite surface protein (anti-CSP), which is extremely abundant on the sporozoite surface, to allow us to find all the sporozoites within the tissue.
- In a midgut with a high oocyst burden, it is not typically challenging to find oocysts using either protein density or nucleic acid stains. By contrast, it can be very difficult to find oocysts in midguts with a low oocyst burden. In these instances, we would recommend use of an oocyst-specific marker, such an anti-capsule antibody for example.
- U-ExM overcomes the impermeability of the oocyst capsule to antibodies and so no specific permeabilization step is required.
- In our experience, images reconstruct dramatically better when using confocal microscopes compared to widefield microscopes. In expanded parasites, the signal is often highly complex, especially for general stains like NHS Ester, WGA or BODIPY ceramide. This complexity can make it dramatically more difficult to discern different parasite structures when more light in captured in the Z-plane.

**Figure 10:**
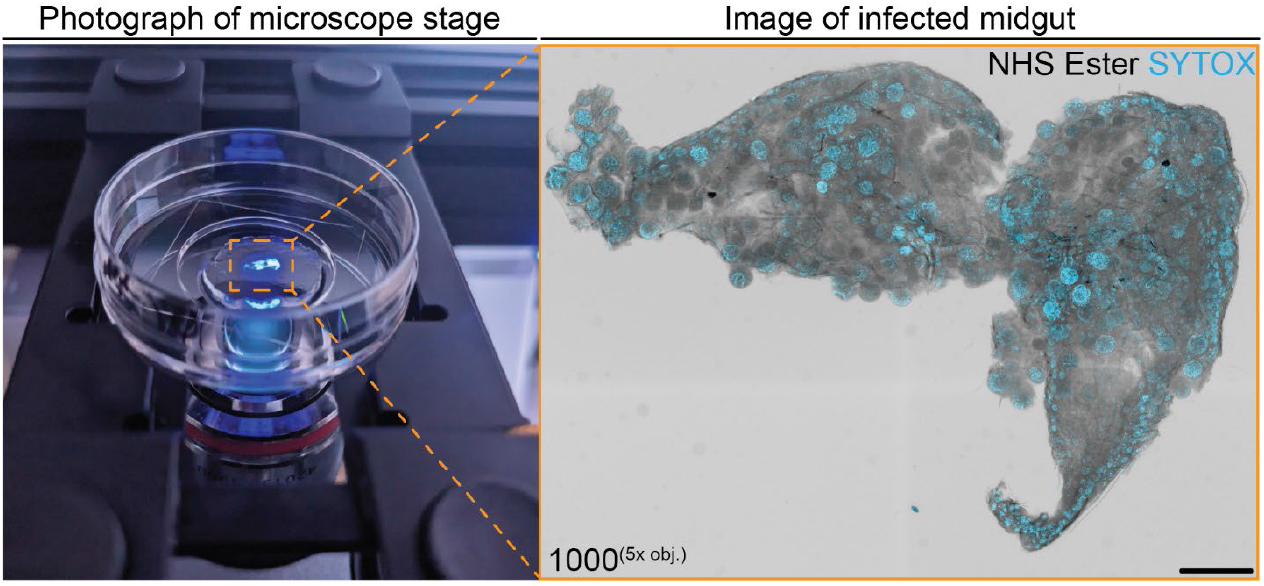
Positioning of expanded mosquito tissues by eye. To aid in the location, positioning, and orientation of infected tissues, they can be visualised by eye using the lamp on a microscope. Pictured left (photograph) is an infected midgut (stained with NHS ester Alexa fluor 405 and SYTOX) illuminated by the UV-lamp of a confocal microscope through a 5x objective lens. Pictured right (orange) is the same midgut following imaging by Airyscan microscopy. Midgut was stained with NHS ester (greyscale, protein density) and SYTOX (cyan, DNA). Number in the bottom corner of indicates z-projection depth in µm. obj.= objective lens. Scale bar = 500 µm.

### Image analysis and presentation tips

- Oocysts are highly complex structures and images of them can contain a lot of detail. This can complicate images and make them difficult to interpret. In many conventional immunofluorescence experiments it is common to either show a single z-slice from a z-stack image, or to generate a maximum intensity projection of the entire image. In our experience, making a maximum intensity projection of an entire expanded oocyst makes the image too complicated to observe any detail, while a single z-slice contains too little detail to infer many biological features. We found that making a projection of a small number of slices that show a region/structure of interest (approximately 1 – 2 µm in z-depth) was most useful for presenting images of expanded oocysts.
- In all experiments during the development of this method, we used the protein density stain NHS ester to provide cellular context as it shows many subcellular structures without the need for specific antibodies. To maximise how much the number of structures this channel could visually show us, we presented it in inverse greyscale. Further, the gamma for this channel was typically presented at 0.45 to allow discernment of dimmer structures.
  ∘ NOTE: changing the gamma of a channel means that fluorescence intensity is no longer linear and this should not be performed when comparing two conditions (like treatment vs control).
  ∘ In our experience, 3D renderings using NHS ester typically do not improve visualisation because it stains all proteinaceous structures.

### Troubleshooting and FAQ

- In this protocol, we suggest placing the gel ‘coverslip side’ face down onto the imaging dish. Mosquito tissues, especially salivary glands, may be suspended in the gel in an orientation that makes them more favourable to image from the ‘parafilm side’ instead. As there is no obvious way of telling this before imaging, we recommend first screening tissues from the ‘coverslip side’ first, but if the tissue is too deep to easily image, place it ‘parafilm side’ down on a new imaging dish and see if the change in orientation helps.
- The commonest issue we encountered with U-ExM reagents is batch-to-batch variation in sodium acrylate. Sodium acrylate purity is essential for U-ExM. We have encountered sodium acrylate that turned either yellow, or reddish-brown when in solution. In both instances, the gels had significantly altered physical properties and did not expand properly. When a new batch of sodium acrylate is received, or a new stock solution is made, test gels that do not contain a biological sample should always be made to ensure that the gels themselves are expanding appropriately.
- **Why isn’t my sample fully expanding?** In our experience developing MoTissU-ExM, the only time when sample expansion factor did not approximately correspond to gel expansion factor was when denaturation had not been performed correctly. For example, when gels were denatured in an oven rather than a heat block, and they were not in direct contact with the heat source, our tissues did not expand.
- **Will antibodies that work by conventional immunofluorescence also work by U-ExM?** In our experience most antibodies work by U-ExM, however, in U-ExM the sample is fully denatured and so antibodies that recognise conformational epitopes are unlikely to work on U-ExM samples. We have found that antibodies that work well by Western blot are more likely to work well by U-ExM than those that work well by immunofluorescence microscopy.
- **What do I do if I have an antibody that hasn’t been used on expanded samples before?** Antibodies on expanded samples often have background fluorescence profiles distinct from unexpanded samples^14^ and so independent antibody validation should always be performed for antibodies that haven’t previously been used on expanded samples. Primary and secondary antibody controls should, along with an antibody dilution series should be run the first time using a new antibody.
- **When during the protocol can I leave my sample(s) overnight?** Fixed tissues can remain fixative or PBS for up to a few weeks (see note). Samples can be left in anchoring solution for approximately 7 days if they are transferred to 4 °C to avoid evaporation. Following denaturation, samples can be left in MilliQ water overnight. Samples in primary antibody can be left for 2-3 days at 4 °C. Samples in secondary antibody can be left overnight. Fully expanded samples in MilliQ water following secondary antibody incubation can be left for approximately one week before imaging.
  ∘ NOTE: Tissues stored in fixative for multiple months may experience issues related to over-fixation including an inability to detect parasite DNA.
- **My gels aren’t getting big enough, are getting too big, or are inconsistently sized**. In our experience, there are many factors that can influence gel size. Most commonly we found that monomer solution more than 2 weeks old, or sodium acrylate stocks more than 6 months old, produced gels of inconsistent size. It has also been shown that small changes in BIS concentration can result in significant changes of gel size and that old APS stocks can slightly alter gel size^32^. Most of these issues can be overcome by keeping track of when stock solutions are made and replacing them accordingly.
- **How long will I be able to stain my fully-expanded stained gel for?** This depends on the antibodies and dyes used to visualise your sample. In our experience, most samples should be highly fluorescent for approximately one week. Some dyes will remain highly fluorescent for much longer than this though, such as NHS Ester and SYTOX.
- **I didn’t finish imaging my sample today, can I go back to it later?** Once a gel has been placed in an imaging dish, it can be re-imaged again for approximately one week. To do this, cover the gel with ∼500 µL of MilliQ water or 0.2 % w/v propyl gallate and store at 4 °C, removing the liquid before imaging.

## Acknowledgements

We thank Prof. Tomoko Ishino (Tokyo Medical & Dental University) for generously providing the *P. berghei* RON11^ctrl^ parasites that were used for some of the infections in this study. This work is supported by an AHA Postdoctoral Fellowship (23POST1011626, BL). We thank the IUSM Pharmacology & Toxicology department for facilitating travel between labs through the Career Development Award. We thank Matt Curtis (PhD) and Michael Hester (PhD) (Zeiss Microscopy, White Plains, NY, USA) for their demonstration of a 40x water-immersion objective, helpful discussions about imaging and their expertise in microscopy optics. We thank Dr. Claire Sayers (University of New South Wales) and Annika Binder (Heidelberg University) for their thoughtful feedback on the manuscript. Anti-CSP antibodies were obtained through BEI Resources, NIAID, NIH.

## Notes

### Competing Interest Statement

The authors have declared no competing interest.

